# Altering expression of a vacuolar iron transporter doubles iron content in white wheat flour

**DOI:** 10.1101/131888

**Authors:** James M. Connorton, Eleanor R. Jones, Ildefonso Rodríguez-Ramiro, Susan Fairweather-Tait, Cristobal Uauy, Janneke Balk

## Abstract

Iron deficiency anaemia is a major global health issue, which has prompted mandatory fortification of cereal products with iron salts or elemental iron in many countries around the world. Rather than post-harvest fortification, biofortification - increasing the intrinsic nutritional quality of crops - is a more sustainable way of alleviating nutrient deficiencies. To identify target genes for biofortification of wheat (*Triticum aestivum*), we functionally characterized homologues of the *Vacuolar Iron Transporter* (*VIT*). The wheat genome contains two *VIT* paralogues, *TaVIT1* and *TaVIT2*, which have different expression patterns, but are both low in the endosperm. TaVIT2, but not TaVIT1, was able to transport iron in a yeast complementation assay. TaVIT2 also transported manganese but not zinc. By over-expressing *TaVIT2* under the control of an endosperm-specific promoter, we achieved a 2-fold increase in iron in white flour fractions, exceeding minimum UK legal fortification levels. The high-iron trait was consistent across independent lines and was stable in the next generation and in two different growth conditions. The single-gene approach impacted minimally on plant growth and was also effective in barley. The anti-nutrient phytate was not increased in white flour from the cisgenic wheat lines, suggesting that food products made from it could contribute to improved iron nutrition.

## Introduction

The essential micronutrients iron and zinc are concentrated in the outer layers of cereal grains, which are commonly removed during food processing, for example in polished rice and in white wheat flour. The low mineral concentration in global staple foods contributes to low iron intake, leading to dietary iron and zinc deficiencies that particularly affect the health of children and women because they have higher physiological requirements. To address this issue, mandatory fortification of wheat flour with iron salts or iron powder has been adopted by 84 countries (http://www.ffinetwork.org/global_progress/index.php). In the Americas, Africa and Asia, iron fortification of flours ranges from 30 to 44 μg/g. In Europe, only the UK has a legal requirement for fortification: white and brown flours must contain at least 16.5 μg/g iron (https://www.food.gov.uk/sites/default/files/multimedia/pdfs/breadflourguide.pdf). All these values are well above the 8 ± 3 μg/g found in white flour from commercial varieties of bread wheat (Eagling *et al.*, 2014a).

Biofortification, or breeding for increased micronutrient content in crops, is a sustainable solution to address mineral deficiencies. However, natural variation for iron content in wheat landraces is limited, in contrast to more genetic variation for zinc (Zhao *et al.*, 2009; Velu *et al.*, 2014). A likely explanation for this difference is that iron homeostasis is more tightly controlled than zinc, because iron in its free form is highly toxic to cells. Modern gene technologies can play an important role, in addition to classical breeding, to achieve biofortification targets, but they require a thorough understanding of the biological processes and knowledge of the genes for uptake, distribution, and accumulation of a particular micronutrient. Following the identification of genes involved in plant iron homeostasis, several have been over-expressed to try and increase iron levels, especially in rice (for reviews see Bashir *et al.*, 2013 and Vasconcelos *et al.*, 2017). They include genes for iron transport (*YSL2*, *IRT1, VIT1*; Schroeder *et al.*, 2013); biosynthesis of the iron chelator nicotianamine (*NAS*) and the iron storage protein ferritin (*FER*). In particular over-expression of *NAS2* (Johnson *et al.*, 2011), or a combination of several genes, has increased iron levels in polished rice more than 6-fold, from 2 μg/g to 15 μg/g in the field (Trijatmiko *et al.*, 2016). In wheat, over-expression of ferritin under the control of an endosperm-specific promoter resulted in 1.6 – 1.8 fold more iron in grains in two independent studies (Borg *et al.*, 2012; Singh *et al.*, 2016). A follow-up study using Synchrotron-based X-Ray Fluorescence showed that iron accumulated in the groove region and remained below detection levels in the endosperm (Neal *et al.*, 2013). Constitutive expression of the rice *NAS2* gene in wheat resulted in 2.1-fold more iron in grains and 2.5-fold more iron in white flour, but combining *NAS2* and *FER* transgenes resulted in only 1.7-fold more iron in grain (Singh *et al.*, 2016). These studies suggest that ferritin is not ideal as a storage sink for iron in cereal grains. Rather, vacuoles may be the dominant store of iron, as demonstrated for Arabidopsis seeds (Kim *et al.*, 2006; Ravet *et al.*, 2009). NanoSIMS studies showed that iron is concentrated in vacuoles in the wheat aleurone (Moore *et al.*, 2012). Therefore we focussed on iron transport into vacuoles as a strategy to biofortify wheat.

## Results

### Wheat has two functionally differentiated *VIT* paralogues

The newly sequenced and annotated wheat genome (International Wheat Genome Sequencing Consortium, 2014; Clavijo *et al.*, 2016) offered the opportunity to make a complete inventory of putative metal transporters in wheat (Borrill *et al.*, 2014). We found that wheat has two *Vacuolar Iron Transporter* genes (*TaVIT1* and *TaVIT2*) on chromosome groups 2 and 5, respectively. Each *TaVIT* gene is represented by 3 copies (homoeologues) from the A, B and D genomes which share 99% identity at the amino acid level (Table S1, Figure S1). TaVIT1 and TaVIT2 have ∼87% amino acid identity with their closest rice homolog, OsVIT1 and OsVIT2, respectively (Zhang *et al.*, 2012). Phylogenetic analysis suggests an early evolutionary divergence of the two *VIT* genes, as there are two distinctly branching clades in the genomes of monocotyledonous species, in contrast to one clade in dicotyledons (Figure 1a). The gene expression profiles of *TaVIT1* and *TaVIT2* were queried across 418 RNA-seq samples (Borrill *et al.*, 2016). All homoeologues of *TaVIT2* were in general more highly expressed than *TaVIT1* homoeologues. In the grains, *TaVIT1* and *TaVIT2* are both expressed in the aleurone, correlating with high levels of iron in this tissue (O’Dell *et al.*, 1972; Moore *et al.*, 2012) which is removed from white flours during the milling process. In contrast, expression of *TaVIT1* and *TaVIT2* is very low in the starchy endosperm, the tissue from which white flour is extracted (Figure 1b). Taken together, differences in phylogeny and expression pattern suggest that TaVIT1 and TaVIT2 may have distinct functions.

**Figure 1.**
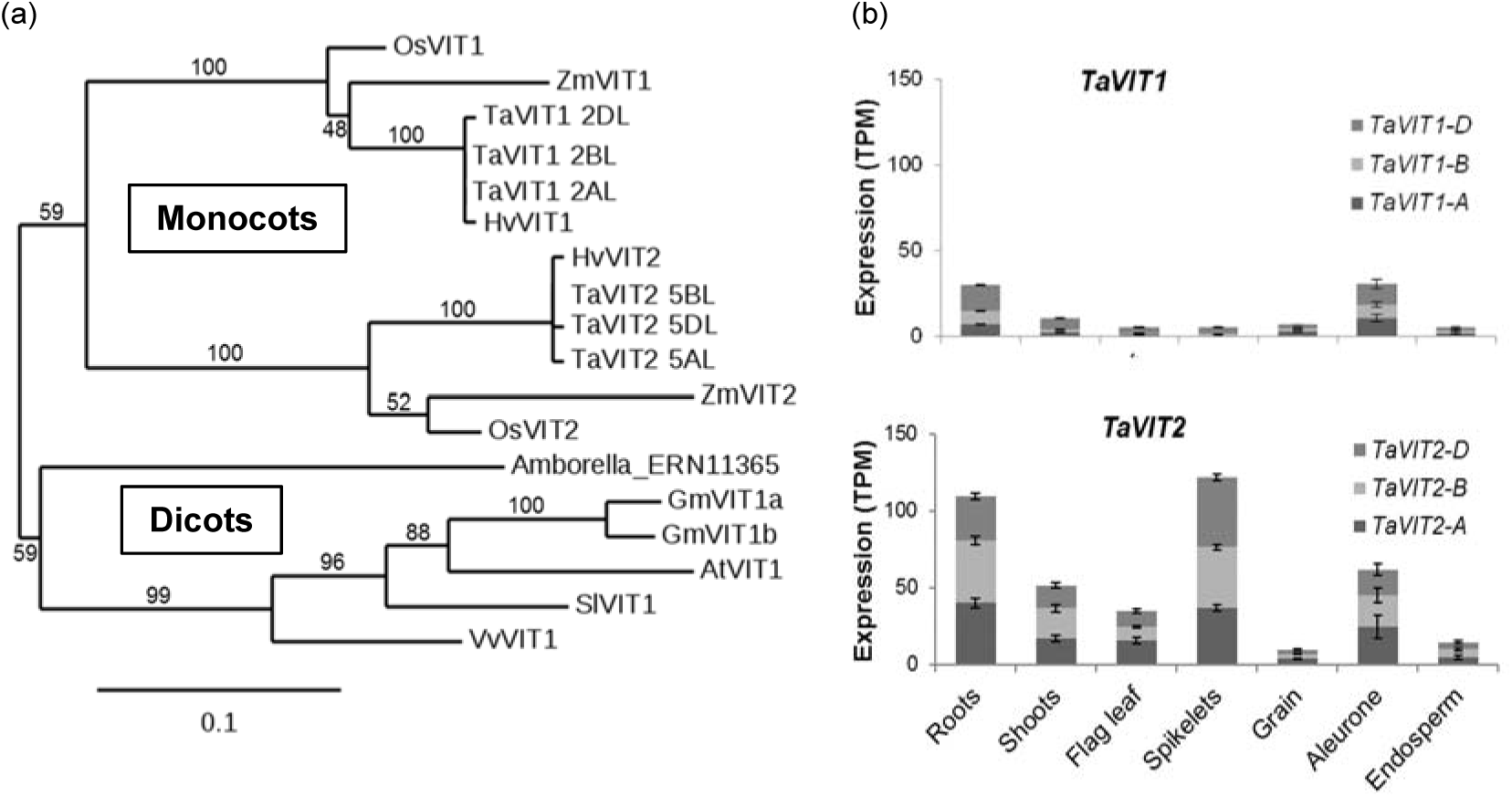
The wheat genome encodes two *VIT* paralogs with different expression patterns. (a) Phylogenetic tree of *VIT* genes from selected plant species: *At, Arabidopsis thaliana*; *Gm, Glycine max* (soybean); *Hv, Hordeum vulgare* (barley); *Os, Oryza sativa* (rice); *Sl, Solanum lycopersicum* (potato); *Ta, Triticum aestivum* (wheat); *Vv, Vitis vinifera* (grape); *Zm, Zea mays* (maize). Numbers above or below branches represent bootstrapping values for 100 replications. (b) Gene expression profiles of *TaVIT1* and *TaVIT2* homoeologs using RNA-seq data from expVIP. Bars indicate mean transcripts per million (TPM) ± SEM, full details and metadata in Table S2.

### TaVIT2 has the ability to transport iron and manganese

To test if the TaVIT proteins transport iron, the 2BL *TaVIT1* homoeolog and 5DL *TaVIT2* homoeolog, hereafter referred to as *TaVIT1* and *TaVIT2* respectively, were expressed in yeast lacking the vacuolar iron transporter Ccc1. The Δ*ccc1* yeast strain is sensitive to high concentrations of iron in the medium because of its inability to store iron in the vacuole. *TaVIT2* fully rescued growth of Δ*ccc1* yeast exposed to a high concentration of FeSO_4_, but *TaVIT1* was no different from the empty vector control (Figure 2a). Yeast Ccc1 can transport both iron and manganese (Lapinskas *et al.*, 1996). Therefore, we carried out yeast complementation using the Δ*pmr1* mutant, which is unable to transport manganese into Golgi vesicles and cannot grow in the presence of toxic levels of this metal (Lapinskas *et al.*, 1995). We found that expression of *TaVIT2* in Δ*pmr1* yeast partially rescued the growth impairment on high concentrations of MnCl_2_, indicating that TaVIT2 can transport manganese (Figure 2b). We also tested if *TaVIT1* and *TaVIT2* were able to rescue growth of the yeast Δ*zrc1* strain, which is defective in vacuolar zinc transport, but neither *TaVIT* gene was able to rescue growth on high zinc (Figure 2c).

**Figure 2.**
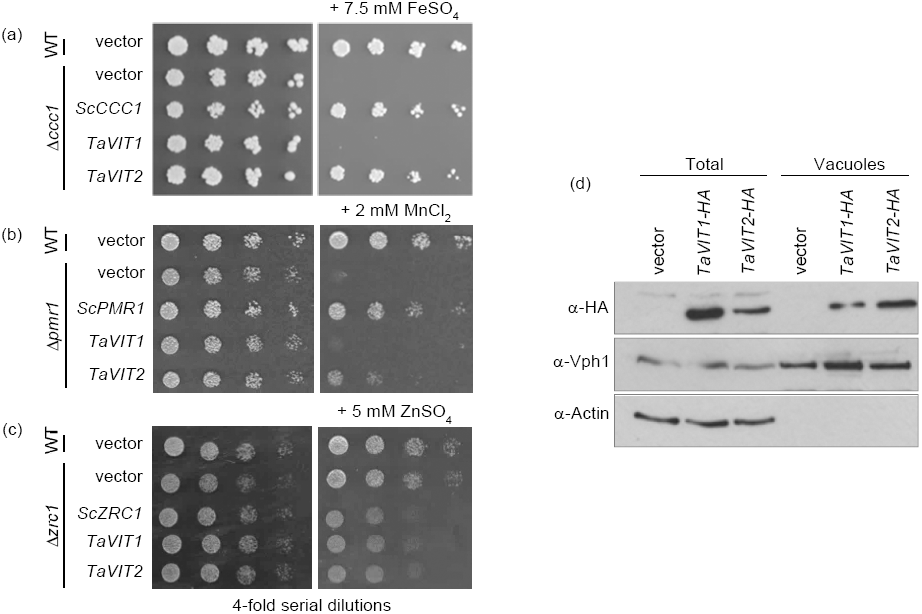
TaVIT2 is a functional iron transporter. (a,b,c) Yeast complementation assays of *TaVIT1* and *TaVIT2* in Δ*ccc1* (a), Δ*pmr1* (b) and Δ*zrc1* (c) compared to yeast that is wild type (WT) for these three genes. The yeast (Sc) *CCC1*, *PMR1* and *ZRC1* genes were used as positive controls. Cells were spotted in a 4-fold dilution series and grown for 2-3 days on plates ± 7.5 mM FeSO_4_ (Δ*ccc1*), 2 mM MnCl_2_ (Δ*pmr1*) or 5 mM ZnSO_4_ (Δ*zrc1*). (d) Immunoblots of total and vacuolar protein fractions from yeast cells expressing haemagglutinin (HA)-tagged *TaVIT1* or *TaVIT2*. The HA-tag did not inhibit the function of TaVIT2 as it was able to complement Δ*ccc1* yeast (data not shown). Vhp1 was used as a vacuolar marker and the absence of actin shows the purity of the vacuolar fraction.

Western blot analysis showed that both proteins were produced in yeast, but that TaVIT1 and TaVIT2 might differ in their intracellular distribution (Figure 2d). TaVIT2 was abundant in vacuolar membranes, co-fractionating with the vacuolar marker protein Vph1. TaVIT1 was also found in the vacuolar membrane fraction, but based on higher abundance in the total fraction, it appeared that most of the TaVIT1 protein was targeted to other membranes. Closer inspection of the amino acid sequences revealed that TaVIT2 contains a universally conserved dileucine motif for targeting to the vacuolar membrane (Bonifacino and Traub, 2003), which is absent from TaVIT1 (Figure S1b).

### Over-expression of *TaVIT2* in the endosperm of wheat specifically increased the iron concentration in white flour

The functional characterization of TaVIT1 and TaVIT2 suggested that TaVIT2, as a *bona fide* iron transporter localized to vacuoles, is a good candidate for iron biofortification. We placed the *TaVIT2* gene under the control of the wheat endosperm-specific promoter of the *High Molecular Weight Glutenin-D1* (*HMW*) gene (Lamacchia *et al.*, 2001) (Figure 3a) and transformed the construct together with a hygromycin resistance marker into the wheat cultivar ‘Fielder’. A total of 27 hygromycin-resistant plants were isolated and the copy number of the transgene was determined by qPCR. There were ten lines with a single copy insertion, and the highest number of insertions was 30. The transgene copy number correlated well with expression of *TaVIT2* in the developing grain (R^2^ = 0.60, *p* < 0.01, Figure S2; Figure 3b). *TaVIT2* expression was increased 3.8 ± 0.2-fold in single copy lines and more than 20-fold in lines with multiple transgenes compared to non-transformed controls.

**Figure 3.**
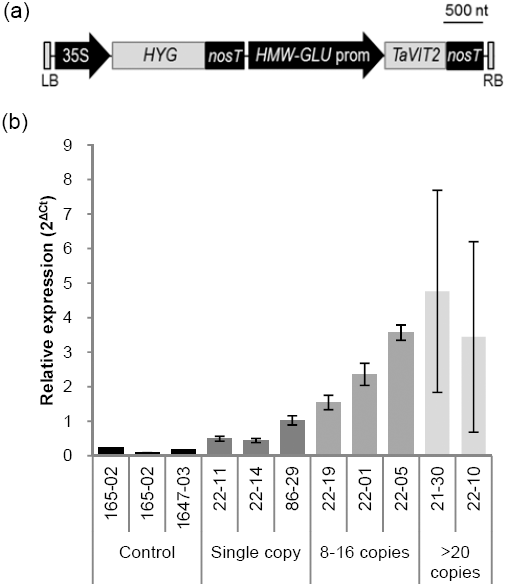
Expression of *TaVIT2* in cisgenic lines. (a) Diagram of the transfer-DNA construct: LB, left border; 35S, CaMV 35S promoter; *HYG,* hygromycin resistance gene; *nosT*, *nos* terminator; *HMW-GLU* prom, high molecular weight glutenin-D1-1 promoter; *TaVIT2*, wheat *VIT2-D* gene; RB, right border. (b) Relative expression levels of *TaVIT2* in developing grains at 10 days post anthesis as determined by quantitative real-time PCR and normalized to housekeeping gene *Traes_4AL_8CEA69D2F.* Plant identification numbers and copy number of the *HMW-TaVIT2* gene are given below the bars. Bars indicate the mean ± SEM of 3 independent biological replicates.

Mature wheat grains from transgenic lines and non-transformed controls were dissected with a platinum-coated blade and stained for iron using Perls’ Prussian Blue. In non-transformed controls, positive blue staining was visible in the embryo, scutellum and aleurone layer (Figure 4), but the endosperm contained little iron. In lines over-expressing *TaVIT2* the Perls’ Prussian Blue staining was visibly increased, in particular around the groove and in patches of the endosperm. To quantify the amount of iron, grains were milled to produce wholemeal flour and a white flour fraction, and analysed by Inductively Coupled Plasma-Optical Emission Spectroscopy (ICP-OES) (Figure 5a, Table S3). Iron levels were consistently enhanced 2-fold in white flour, from 9.7 ± 0.3 μg/g in control lines to 21.7 ± 2.7 μg/g in lines with a single copy of *HMW*-*TaVIT2* (*p* < 0.001). Additional transgene copies resulted in a similar 2-fold increase in iron, whereas lines with ≥ 20 copies contained 4-fold more iron than controls, to 41.5 ± 8.2 μg/g in white flour (*p* < 0.001). The iron content of wholemeal flour of single insertion lines was similar to control lines, but increased up to 2-fold in high copy lines. No major differences were found for other mineral nutrients in single-copy *HMW*-*TaVIT2* wheat grains, such as zinc, manganese and magnesium (Table S3), nor for the heavy metal contaminants cadmium and lead (Table S4).

**Figure 4.**
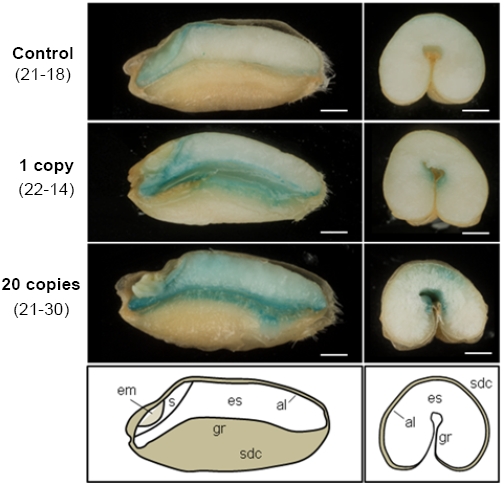
Perls’ Prussian Blue staining for iron in grains transformed with *HMW-TaVIT2*. Grains from T_0_ wheat plants were dissected longitudinally (left) or transversely (right). em, embryo; s, scutellum; sdc, seed coat; es, endosperm; al, aleurone, gr, groove. The transgene copy number and line number are indicated on the far left. Scale bars = 1 mm.

**Figure 5.**
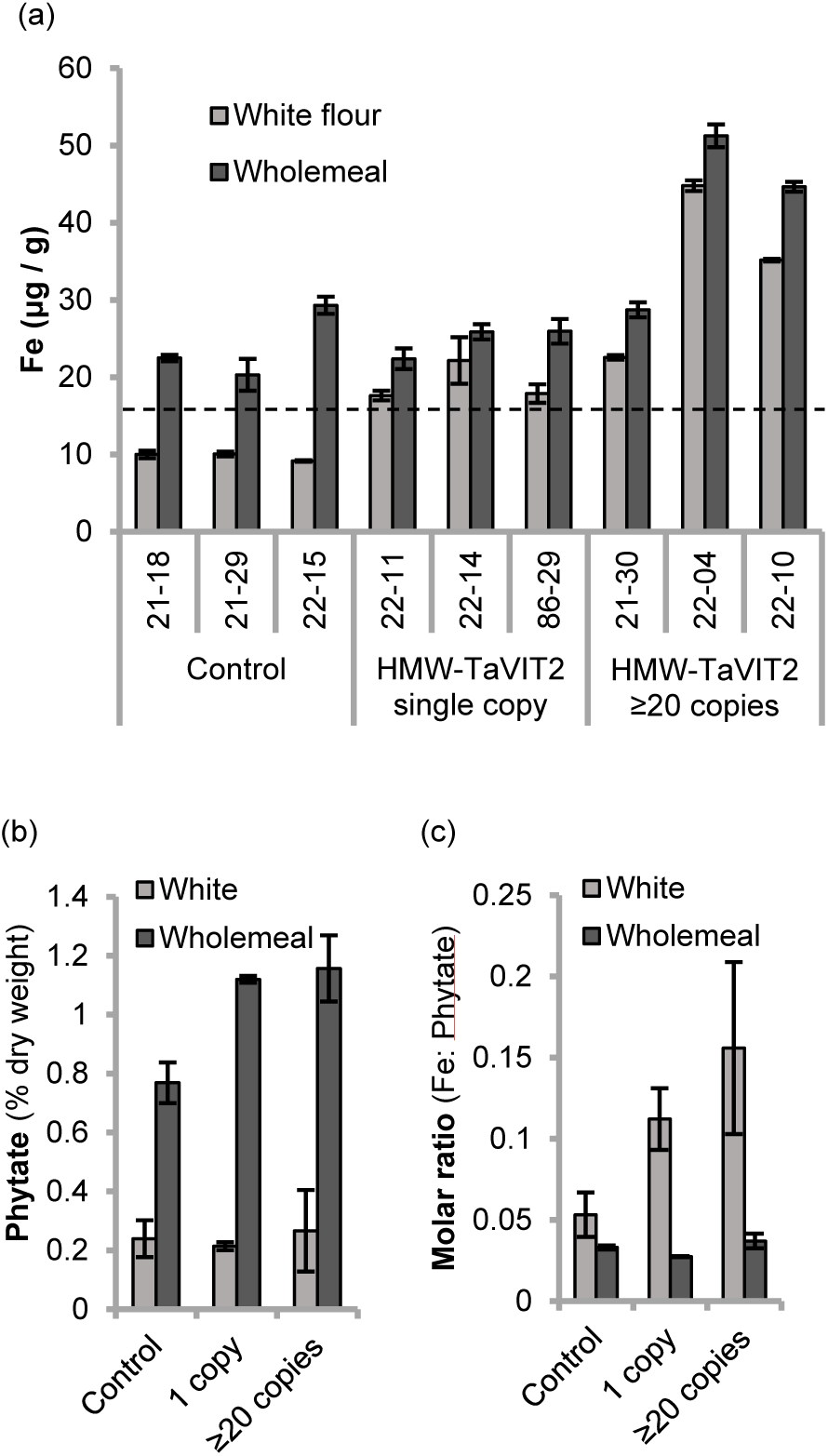
Iron and phytate content of flour milled from *HMW-TaVIT2* wheat lines. (a) Iron concentrations in white and wholemeal flour from 3 control and 6 *HMW-TaVIT2* lines. Bars represent the mean of 2 technical replicates and the deviation of the mean. White flour from *HMW-TaVIT2* lines has significantly more iron than control lines (n = 3-4, p<0.001; see Table S3 for all data). The dotted line at 16.5 μg/g iron indicates the minimum requirement for wheat flour sold in the UK. (b) Phytate content of white and wholemeal flour of control and *HMW-TaVIT2* expressing wheat. Bars represent the mean of 2 biological replicates ± deviation of the mean. (c) Molar ratio of iron:phytate in control and *HMW-TaVIT2* expressing lines. Bars represent the mean of 2 biological replicates and the deviation of the mean.

### The iron:phytate ratio is improved in white flour

Iron in plants is almost exclusively in the form of non-heme iron, and bioavailability of dietary iron may be as low as 5-10% in populations consuming monotonous plant-based diets (Zimmermann and Hurrell, 2007). The absorption of non-heme iron is increased by meat and ascorbic acid, but inhibited by phytates, polyphenols, and, to a lesser extent, calcium (Hurrell and Egli, 2010). Cereal grains contain high levels of phytate (myo-inositol-1,2,3,4,5,6-hexakisphosphate), a phosphate storage molecule for the plant, which is concentrated in the aleurone tissue of rice and wheat, but is low in the endosperm (O’Dell *et al.*, 1972). We measured phytate levels in white flour of the *TaVIT2* over-expressing lines, but found no significant increase in phytate or total phosphorus (Figure 5b; Table S3). There was a small increase in phytate in wholemeal flour produced from the *HMW-TaVIT2* lines. Considering the 2-fold increase in iron, the iron:phytate molar ratio was improved 2-fold in white flour of *HMW-TaVIT2* lines, and there was no change in the ratio in wholemeal flour (Figure 5c). It was previously shown that iron in bread made from white flour is taken up by Caco-2 cells, in a widely-used intestinal cell culture model, but iron in wholemeal bread was not (Eagling *et al.*, 2014b). Therefore, relocating iron to the endosperm may be more effective than increasing total iron in the grain as a biofortification strategy.

### Endosperm-specific over-expression of *TaVIT2* does not affect growth parameters

To investigate if *TaVIT2* over-expression affected plant growth, we measured plant height, tiller number, number of grains per plant and thousand-grain weight in *TaVIT2* over-expressing lines and controls. None of these growth parameters were negatively affected by the *HMW*-*TaVIT2* transgene in the T_0_ generation grown in controlled environment rooms (Figure 6 and Table S5). The high-iron trait was stable in the next generation (T_1_), co-segregating in a 3:1 ratio (χ^2^=0.29) with the presence of the *HMW-TaVIT2* transgene (Table S6). Further greenhouse experiments confirmed no significant differences in growth and yield component traits for *HMW-TaVIT2* plants compared to wild-type segregants or controls.

**Figure 6.**
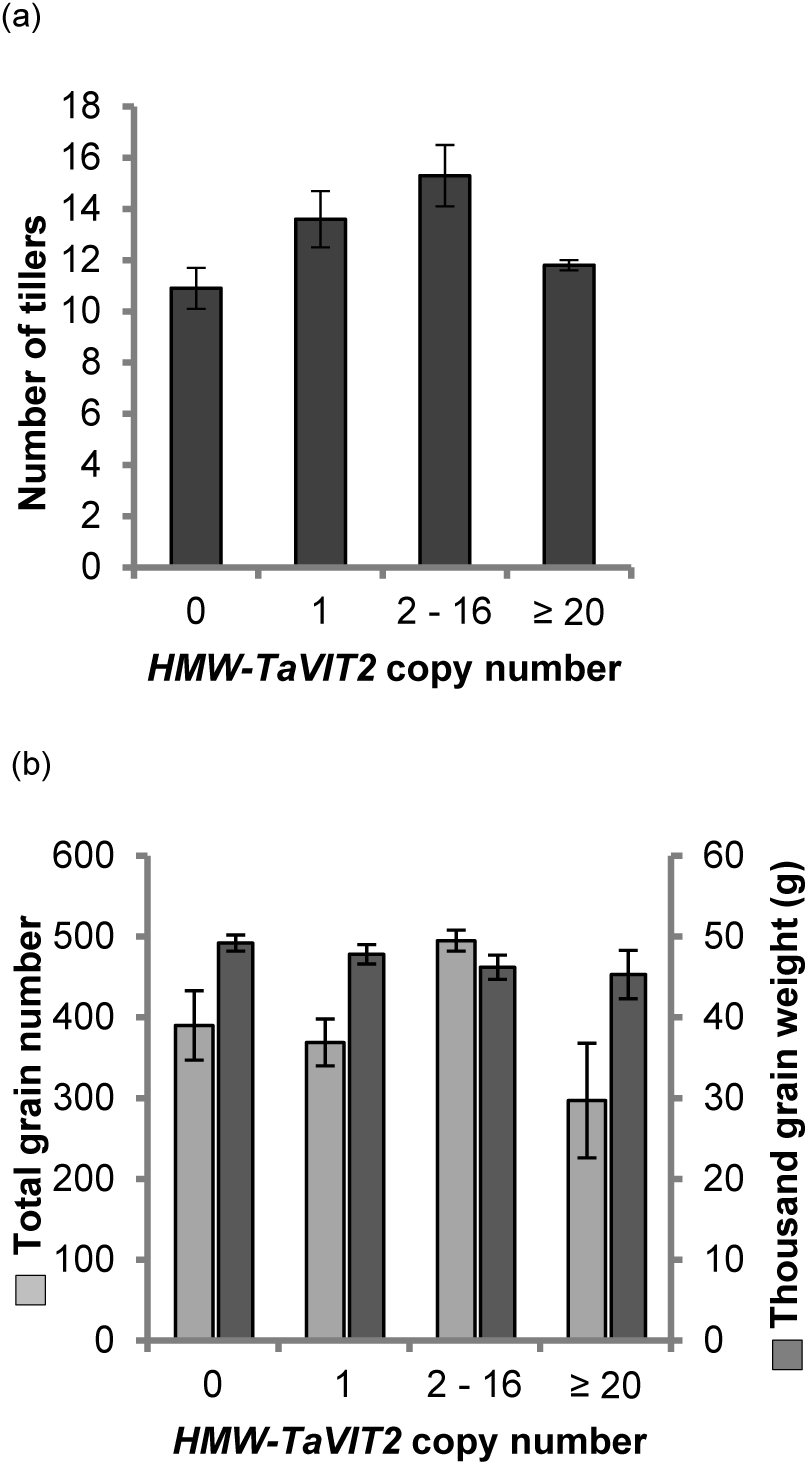
Growth parameters of *HMW-TaVIT2* wheat. (a) Number of tillers and (b) seed output of T_0_ wheat plants with indicated *HMW-TaVIT2* copy numbers. Bars indicate mean ± SEM of the following numbers of biological replicates: zero gene copies, n = 9; 1 gene copy, n = 10; 2 −16 gene copies, n = 9; ≥ 20 gene copies, n = 6. Further details given in Table S5.

### Expression of *HMW*-*TaVIT2* in barley increased grain iron and manganese content

We also transformed barley (*Hordeum vulgare* cv. Golden Promise) with the *HMW*-*TaVIT2* construct. The 12 transgenic plants had either 1 or 2 copies of the transgene and were indistinguishable from non-transformed controls with regards to vegetative growth and grain development. Lines B2 (1 copy) and B3 (2 copies) were selected for analysis and found to contain 2-fold more iron than the control in both white and wholemeal flour (Figure 7). Interestingly, in barley there was also a 2-fold increase in manganese levels (Figure 7), which could be expected from the ability of TaVIT2 to transport manganese in yeast (Figure 2b). Together, our results indicate that endosperm-specific over-expression of*TaVIT2* is a successful strategy for increasing the iron content in different cereal crop species.

**Figure 7.**
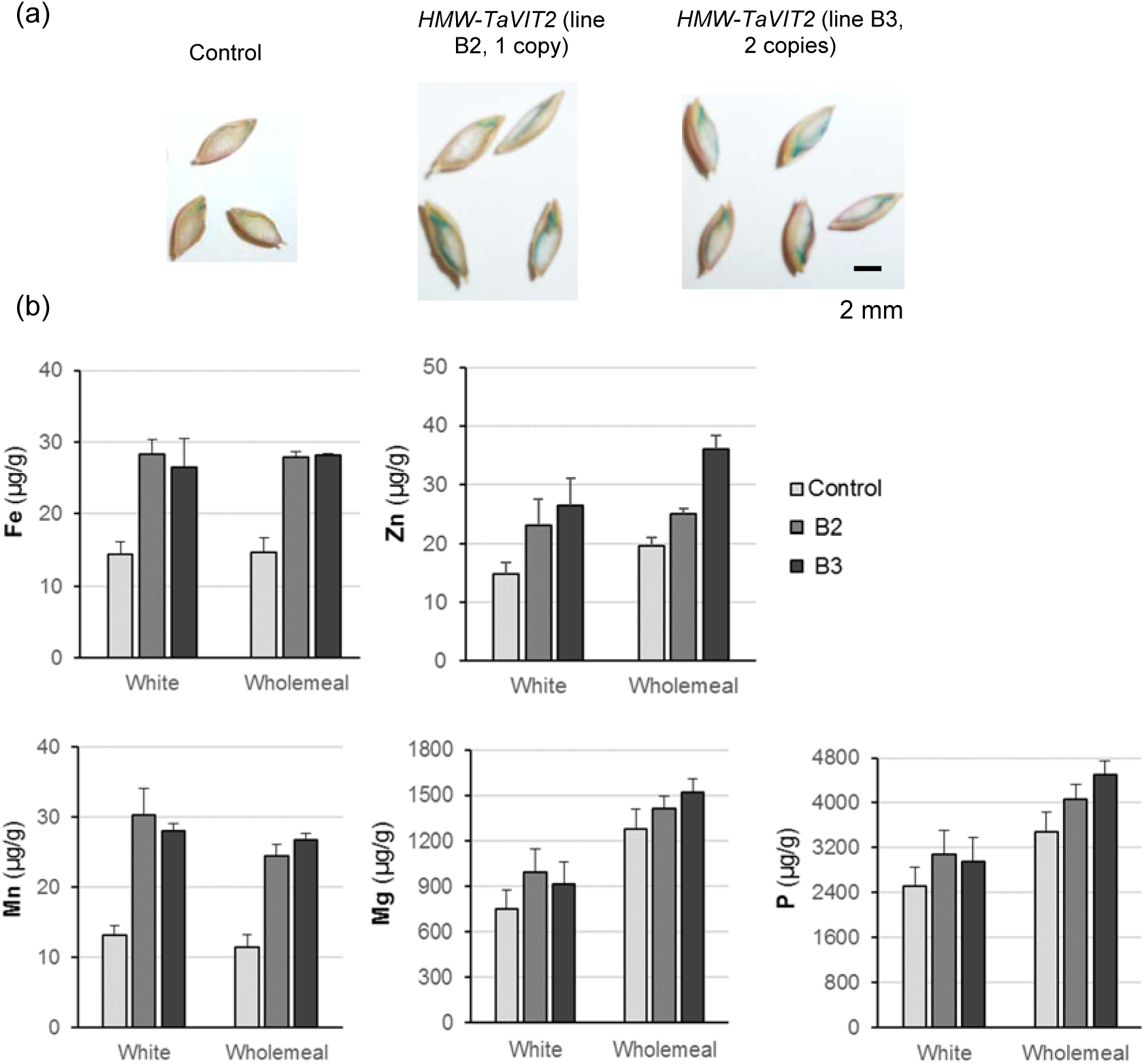
Endosperm-specific over-expression of *TaVIT2* in barley. The *TaVIT2-5DL* gene from wheat under the control of the wheat *HMW-GLU-1D-1* promoter (see Figure 3a for full details) was transformed into barley (*Hordeum vulgare* var. Golden Promise). Positive transformants were selected by hygromycin. (a) Mature barley grains of a control plant and two transgenic lines stained with Perls’ Prussian blue staining for iron. (b) Element analysis in white and wholemeal flours from a control and two *HMW-TaVIT2* over-expressing barley plants. The values are the mean of 2 technical replicates. Error bars represent the deviation of the mean.

## Discussion

In this study we showed that the recently sequenced wheat genome greatly facilitates the design of cisgenic approaches for biofortification in this important staple crop. Over-expression of the vacuolar iron transporter *TaVIT2* in wheat endosperm was very effective in raising the iron concentration in this tissue. A similar approach, but using the *VIT1* gene from *Arabidopsis thaliana* under the control of a tuber-specific promoter, has been shown to dramatically increase iron in cassava tubers (Narayanan *et al.*, 2015). We hypothesize that increased sequestration of iron in the vacuoles creates a sink which then upregulates the relocation of iron to that tissue. If the tissue normally stores iron in vacuoles rather than in ferritin, proteins and chelating molecules for iron mobilisation into the vacuole will already be present. For a sink-driven strategy, timely expression of the gene in a specific tissue is essential: if a sink is expressed constitutively, for example using the CaMV 35S promoter, then it is unlikely to be effective in directing iron to a particular tissue. Interestingly, knock-out mutants of *VIT1* and *VIT2* in rice accumulated more iron in the embryo (Zhang *et al.*, 2012). A likely scenario is that iron distributed to the developing rice grain cannot enter the vacuoles in the aleurone, and is thus diverted to the embryo. The finding further supports the idea that VITs play a key role in iron distribution in cereal grains. Another advantage of endosperm-specific expression is that possible growth defects in vegetative tissues are avoided.

Wheat and barley transformed with the same *HMW-TaVIT2* construct showed surprising differences in the accumulation of iron and manganese. Wheat had a 2-fold increase in iron in the endosperm only, whereas barley contained 2-fold more iron in whole grains. Barley grains also contained 2-fold more manganese, but this element was not increased in wheat, even though TaVIT2 was found to transport both iron and manganese in yeast complementation assays. It is possible that the wheat *HMW* promoter has a different expression pattern in barley. If the promoter is activated in the aleurone cells in addition to the endosperm, this may lead to the observed higher iron concentrations in whole barley grains. The pattern of promoter activity can be further investigated with reporter constructs or by in-situ hybridization specific for the transgene. It is also possible that wheat and barley differ in iron and manganese transport efficiency from roots to shoots, thus affecting the total amount of iron and manganese that is (re)mobilized to the grain.

While over-expression of *TaVIT2* as a single gene approach resulted in a robust 2-fold increase in iron, combining this with constitutive *NAS* over-expression may increase the iron in white flour even further. Nicotianamine is also reported to improve the bioavailability of iron (Lee *et al*., 2009), which is a major determinant for the success of any biofortification strategy. The bioavailability of iron in bread made from *HMW-TaVIT2* lines will be addressed in future studies, using the well-established in vitro assay with Caco-2 cells or, in the longer term, human studies. Another major factor for biofortified wheat will be the acceptance by consumers. We hope that a cisgenic strategy will help to swing public opinion towards highly beneficial applications of modern gene technology. The vector sequences and hygromycin marker can be removed with CRISPR technology, leaving only wheat DNA. In addition, better understanding of mineral translocation into the endosperm, using *VIT2* over-expression as proof-of-principle, may help to identify other gene targets for biofortification that can be targeted by non-transgenic approaches such as TILLING (Krasileva *et al.*, 2017).

## Experimental procedures

### Identification of wheat *VIT* genes, phylogenetic analysis and analysis of RNA-seq data

The coding sequences of the wheat *VIT* genes were found by a BLAST search of the rice *OsVIT1 (LOC_Os09g23300)* and *OsVIT2 (LOC_Os04g38940)* sequences in Ensembl Plants (http://plants.ensembl.org). Full details of the wheat genes are given in Table S1. Sequences of *VIT* genes from other species were found by a BLAST search of the Arabidopsis *AtVIT1 (AT2G01770)* and rice *VIT* sequences against the Ensembl Plants database. Amino acid alignments were performed using Clustal Omega (http://www.ebi.ac.uk/Tools/msa/clustalo/). The tree was plotted with BioNJ with the Jones-Taylor-Thornton matrix and rendered using TreeDyn 198.3. RNA-seq data was obtained from the expVIP database (Borrill *et al.*, 2016; http://www.wheat-expression.com). Full details of the data-sets used are given in Table S2.

### Yeast complementation

Coding DNA sequences for the wheat 2BL *VIT1* homoeolog (TRIAE_CS42_2BL_TGACv1_129586_AA0389520) and the 5DL *VIT2* homoeolog (TRIAE_CS42_5DL_TGACv1_433496_AA1414720) were synthesised and inserted into pUC57 vectors by Genscript (Piscataway, NJ, USA). The wheat *VIT* genes were first synthesised with wheat codon usage, but *TaVIT1* was poorly translated in yeast so was re-synthesized with yeast codon usage including a 3x haemagglutinin (HA) tag at the C-terminal end. Untagged codon-optimised *TaVIT1* was amplified from this construct using primers *TaVIT1co-Xba*I-F and *TaVIT1co-EcoR*I-R (see Table S7 for primer sequences). *TaVIT2-HA* was cloned by amplifying the codon sequence without stop codon using primers *TaVIT2-BamH*I-F and *TaVIT2*(ns)-*EcoR*I-R, and by amplifying the HA tag using primers HAT-*EcoR*I-F and HAT(Stop)-*Cla*I-R. The two DNA fragments were inserted into plasmid p416 behind the yeast *MET25* promoter (Mumberg *et al.*, 1995).

Genes *ScCCC1* and *ScZRC1* were cloned from yeast genomic DNA, using the primer pairs *ScCCC1*-*BamH*I-F and *ScCCC1*-*EcoR*I-R, and *ScZRC1*-*Xba*I-F and*ScZRC1*-*EcoR*I-R, respectively. Following restriction digests the DNA fragments were ligated into vector p416-*MET25* and confirmed by sequencing. All constructs were checked by DNA sequencing.

The *Saccharomyces cerevisiae* strain BY4741 (MATa *his3*Δ1 *leu2*Δ0 *met15*Δ0 *ura3*Δ0) was used in all yeast experiments. Either wild-type (WT), Δ*ccc1* (Li *et al.*, 2001), Δ*zrc1* (MacDiarmid *et al.*, 2003) or Δ*pmr1* (Lapinskas *et al.*, 1995) was transformed with approximately 100 ng DNA using the PEG/lithium acetate method (Ito *et al.*, 1983). Complementation analysis was performed via drop assays using overnight cultures of yeast grown in selective synthetic dextrose (SD) media, diluted to approximately 1 × 10^6^ cells/ml, spotted in successive 4 × dilutions onto SD plates containing appropriate supplements. Plates were incubated for 3 days at 30°C. Total yeast protein extraction was performed by alkaline lysis of overnight cultures (Kushnirov, 2000).

### Preparation of vacuoles from yeast

Preparation of yeast vacuoles was performed using cell fractionation over a sucrose gradient (Nakanishi *et al.*, 2001; Hwang *et al.*, 2000). Briefly, 1 L yeast was grown in selective SD media to an OD_600_ of 1.5-2.0 then centrifigued at 4000 ***g*** for 10 min, washed in buffer 1 (0.1 M Tris-HCl pH 9.4, 50 mM β-mercaptoethanol, 0.1 M glucose) and resuspended in buffer 2 (0.9 M sorbitol, 0.1 M glucose, 50 mM Tris-2-(N-morpholino)ethanesulfonic acid (MES) pH 7.6, 5 mM dithiothreitol (DTT), 0.5 × SD media). Zymolyase 20T (Seikagaku, Tokyo, Japan) was added at a concentration of 0.05% (w/v) and cells were incubated for 2 h at 30°C with gentle shaking. After cell wall digestion, spheroplasts were centrifuged at 3000 ***g*** for 10 min and then washed in 1 M sorbitol before being resuspended in buffer 3 (40 mM Tris-MES, pH 7.6, 1.1 M glycerol, 1.5% (w/v) polyvinylpyrrolidone 40,000; 5 mM EGTA, 1 mM DTT, 0.2% (w/v) bovine serum albumin (BSA), 1 mM phenylmethylsulfonyl fluoride (PMSF), 1 × protease inhibitor cocktail (Promega)) and homogenised on ice using a glass homogeniser. The homogenate was centrifuged at 2000 ***g*** for 10 min at 4°C and the supernatant was transferred to fresh tubes, while the pellet was resuspended in fresh buffer 3 and centrifuged again. The supernatants were pooled and centrifuged at 150,000 ***g*** for 45 min at 4°C to pellet microsomal membranes. For preparation of vacuole-enriched vesicles the pellet was resuspended in 15% (w/w) sucrose in buffer 4 (10 mM Tris-MES pH7.6, 1 mM EGTA, 2 mM DTT, 25 mM KCl, 1.1 M glycerol, 0.2% (w/v) BSA, 1 mM PMSF, 1 × protease inhibitor cocktail) and this was layered onto an equal volume of 35% (w/w) sucrose solution in buffer 4 before centrifugation at 150,000 ***g*** for 2 h at 4°C. Vesicles were collected from the interface and diluted in buffer 5 (5 mM Tris-MES pH 7.6, 0.3 M sorbitol, 1 mM DTT, 1 mM EGTA, 0.1 M KCl, 5 mM MgCl_2,_ 1 mM PMSF, 1 × protease inhibitor cocktail). The membranes were centrifuged at 150,000 ***g*** for 45 min at 4°C and resuspended in a minimal volume of buffer 6 (5 mM Tris-MES pH 7.6, 0.3 M sorbitol, 1 mM DTT, 1 mM PMSF, 1 × protease inhibitor cocktail). Vesicles were snap-frozen in liquid nitrogen and stored at −80°C.

### Generation of transgenic plant lines

The *TaVIT2* gene was amplified using primers TaVIT2-*Nco*IF and TaVIT2-*Spe*IR and cloned into vector pRRes14_RR.301 containing the promoter sequence comprising nucleotides −1187 to −3 with respect to the ATG start codon of the *GLU-1D-1* gene, which encodes the high-molecular-weight glutenin subunit 1Dx5 (Lamacchia *et al.*, 2001). The promoter-gene fragment was then cloned into vector pBract202 containing a hygromycin resistance gene and LB and RB elements for insertion into the plant genome (Smedley and Harwood, 2015). The construct was checked by DNA sequencing. Transformation into wheat (cultivar Fielder) and barley (cultivar Golden Promise) were performed by the BRACT platform at the John Innes Centre using *Agrobacterium*-mediated techniques as described previously (Wu *et al.*, 2003; Harwood *et al.*, 2009). Transgene insertion and copy number in T_0_ plants were assessed by iDNA Genetics (Norwich, UK) using qPCR with a Taqman probe. For the T_1_ generation, the presence of the hygromycin resistance gene was analysed by PCR with primers Hyg-F and Hyg-R.

### Plant growth and quantitative analysis

The first generation of transgenic plants (T_0_) were grown in a controlled environment room under 16 h light (300 μmol m^-2^ s^-1^) at 20°C / 8 h dark at 15°C with 70% relative humidity. The next generation (T_1_) were grown in a glasshouse kept at approximately 20°C with 16 h light. Ears from wheat and barley plants were threshed by hand and grain morphometric characteristics, mass and number were determined using a MARVIN universal grain analyser (GTA Sensorik, GmbH, Neubrandenburg, Germany).

### RNA extraction and qRT-PCR

Samples of developing grain were taken at 10 days post anthesis and frozen in liquid nitrogen. RNA extraction was performed using phenol/chloroform extraction (Box *et al.*, 2011). Developing grains were ground with a pestle and mortar under liquid nitrogen and mixed with RNA extraction buffer (0.1 M Tris-HCl, pH 8; 5 mM EDTA; 0.1 M NaCl, 0.5% (w/v) SDS, 1% (v/v) 2-mercaptoethanol) and Ambion Plant RNA Isolation Aid (ThermoFisher). Samples were centrifuged for 10 min at 15,000 ***g*** and the supernatant was added to 1:1 acidic phenol (pH 4.3):chloroform. After mixing and incubation at room temperature for 10 min, the upper phase was added to isopropanol containing 0.3 M sodium acetate. Samples were incubated at −80°C for 15 min and centrifuged for 30 min at 15,000 ***g*** at 4°C. The supernatant was discarded and the pellet was washed twice in 70% (v/v) ethanol and dried, before being resuspended in RNAse-free water. RNA was DNase treated using TURBO DNase-free kit (ThermoFisher) as per manufacturer’s instructions, DNase inactivation reagent was added and the samples were centrifuged at 10,000 ***g*** for 90 s. Supernatant containing RNA was retained. RNA was reverse transcribed using oligo dT primer and Superscript II reverse transcriptase (ThermoFisher) according to manufacturer’s instructions. Quantitative real time PCR was used to analyse expression of *TaVIT2* and the housekeeping gene (*HKG*) *Traes_4AL_8CEA69D2F*, chosen because it was shown to be the most stable gene expression across grain development in over 400 RNAseq samples (Borrill *et al.*, 2016), using primer pairs qRT-*TaVIT2*-F, qRT-*TaVIT2*-R and qRT-*HKG*-F, qRT-*HKG*-R, respectively. Samples were run in a CFX96 Real-Time System (Bio-Rad) with the following conditions: 3 min at 95°C, 35 cycles of (5 s at 95°C, 10 s at 62°C, 7 s at 72°C), melt curve of 5 s at 65°C and 5 s at 95°C. *TaVIT2* expression levels were normalized to expression levels of the housekeeping gene and expressed as 2^ΔCt^.

### Perls’ Prussian Blue staining

Mature grains were dissected using a platinum-coated scalpel and stained for 45 mins in Perls’ Prussian blue staining solution (2% (w/v) potassium hexacyanoferate (II); 2% (v/v) hydrochloric acid), then washed twice in deionised water.

### Flour preparation, element analysis and phytate determination

Barley grains were de-hulled by hand and all grains were coarsely milled using a coffee grinder then ground into flour using a pestle and mortar. White flour fractions were obtained by passing the material through a 150 μm nylon mesh. Flour samples were dried overnight at 55°C and then digested for 1 h at 95°C in ultrapure nitric acid (55% v/v) and hydrogen peroxide (6% v/v). Samples were diluted 1:11 in ultrapure water and analysed by Inductively Coupled Plasma-Optical Emission Spectroscopy (Vista-pro CCD Simultaneous ICP-OES, Agilent, Santa Clara, CA, USA) calibrated with standards; Zn, Fe and Mg at 0.2, 0.4, 0.6, 0.8 and 1mg/l, Mn and P at 1, 2, 3, 4 and 5 mg/l. Soft winter wheat flour was used as reference material (RM 8438, National Institute of Standards and Technology, USA) and analysed in parallel with all experimental samples. Phytate levels were determined using a phytic acid (total phosphorus) assay kit (Megazyme, Bray, Ireland).

### Statistical analysis

Statistical analyses (F-test, ANOVA with Tukey post-hoc test, Student’s *t*-test, regression analysis, *χ*^2^) were performed using Microsoft Excel 2010 and Genstat 18^th^ Edition. When representative images are shown, the experiment was repeated at least 3 times with similar results.

## Acknowledgements

We would like to thank James Simmonds for technical assistance; Wendy Harwood for wheat transformation; Alison Huttly at Rothamsted Research for the *HMW GLU-D1-1* promoter; Sylvaine Bruggraber at MRC-HNR and Graham Chilvers at UEA for element analysis; Tony Miller and Dale Sanders for helpful discussions. This work was funded by HarvestPlus (J.M.C., J.B. and C.U.) and by the Biotechnology and Biological Sciences Research Council (BBSRC) Institute Strategic Programme Grant BB/J004561/1 (C.U. and J.B.) and DRINC2 grant BB/L025515/1 (I.R.R., S.F.T. and J.B.).

### Author contributions

J.M.C. and E.R.J. designed and performed experiments; All authors analysed and interpreted data; C.U. and J.B. conceived and designed the project; J.M.C. and J.B. co-wrote the paper.

### Conflict of interest

Authors declare no conflict of interest.

## Supporting Information

Addition Supporting information on the following pages:

**Figure S1** Gene models and protein sequence of wheat vacuolar iron transporters.

**Figure S2** Correlation between *HMW-TaVIT2* transgene copy number and expression of *TaVIT2.*

**Table S1** Wheat *VIT* genes identified in this study.

**Table S2** Expression analysis of *TaVIT* genes.

**Table S3** Element analysis of control and *HMW-TaVIT2* wheat lines.

**Table S4** Heavy metals in control and *HMW-TaVIT2* wheat lines.

**Table S5** Architectural and yield components of control and *HMW-TaVIT2* T_0_ transformants.

**Table S6** Architectural and yield components of T_1_ plants segregating from a *TaVIT2* over-expressor T_0_ plant.

**Table S7** List of primers.

## References

Bashir, K., Nozoye, T., Ishimaru, Y., Nakanishi, H., and Nishizawa, N.K. (2013) Exploiting new tools for iron bio-fortification of rice. Biotechnol. Adv. 31, 1624–1633.

Bonifacino, J.S. and Traub, L.M. (2003) Signals for sorting of transmembrane proteins to endosomes and lysosomes. Annu. Rev. Biochem. 72, 395–447.

Borg, S., Brinch-Pedersen, H., Tauris, B., Madsen, L.H., Darbani, B., Noeparvar, S., and Holm, P.B. (2012) Wheat ferritins: Improving the iron content of the wheat grain. J. Cereal Sci. 56, 204–213.

Borrill, P., Connorton, J.M., Balk, J., Miller, A.J., Sanders, D., and Uauy, C. (2014) Biofortification of wheat grain with iron and zinc: integrating novel genomic resources and knowledge from model crops. Front. Plant Sci. 5, 1–8.

Borrill, P., Ramirez-Gonzalez, R., and Uauy, C. (2016) expVIP: a Customizable RNA-seq Data Analysis and Visualization Platform. Plant Physiol. 170, 2172–2186.

Box, M.S., Coustham, V., Dean, C., and Mylne, J.S. (2011) Protocol: A simple phenol-based method for 96-well extraction of high quality RNA from Arabidopsis. Plant Methods 7, 7.

Clavijo, B.J. et al. (2016) An improved assembly and annotation of the allohexaploid wheat genome identifies complete families of agronomic genes and provides genomic evidence for chromosomal translocations. bioRxiv Jan 1, 80796.

Eagling, T., Neal, A.L., McGrath, S.P., Fairweather-Tait, S., Shewry, P.R., and Zhao, F.J. (2014a) Distribution and speciation of iron and zinc in grain of two wheat genotypes. J. Agric. Food Chem. 62, 708–716.

Eagling, T., Wawer, A.A., Shewry, P.R., Zhao, F., and Fairweather-tait, S.J. (2014b) Iron Bioavailability in Two Commercial Cultivars of Wheat: Comparison between Wholegrain and White Flour and the Effects of Nicotianamine and 2'-Deoxymugineic Acid on Iron Uptake into Caco-2 Cells. J. Agric. Food Chem. 62, 10320–10325.

Harwood, W.A., Bartlett, J.G., Alves, S.C., Perry, M., Smedley, M.A., Leyland, N., and Snape, J.W. (2009) Barley transformation using Agrobacterium-mediated techniques. In Methods in Molecular Biology, Transgenic Wheat, Barley and Oats, H.D. Jones and P.R. Shewry, eds (Humana Press), pp. 137–147.

Hwang, I., Harper, J.F., Liang, F., and Sze, H. (2000) Calmodulin activation of an endoplasmic reticulum-located calcium pump involves an interaction with the N-terminal autoinhibitory domain. Plant Physiol. 122, 157–68.

Hurrell, R. and Egli, I. (2010) Iron bioavailability and dietary reference values. Am. J. Clin. Nutr. 91, 1461S–1467S.

International Wheat Genome Sequencing Consortium, (IWGSC) (2014) A chromosome-based draft sequence of the hexaploid bread wheat (*Triticum aestivum*) genome. Science 345, 1251788.

Ito, H., Fukuda, Y., and Murata, K. (1983) Transformation of intact yeast cells treated with alkali cations. J. Bacteriol. 153, 163–168.

Johnson, A.A.T., Kyriacou, B., Callahan, D.L., Carruthers, L., Stangoulis, J., Lombi, E., and Tester, M. (2011) Constitutive overexpression of the *OsNAS* gene family reveals single-gene strategies for effective iron- and zinc-biofortification of rice endosperm. PLoS One 6, e24476.

Kim, S. a, Punshon, T., Lanzirotti, A., Li, L., Alonso, J.M., Ecker, J.R., Kaplan, J., and Guerinot, M. Lou (2006) Localization of iron in Arabidopsis seed requires the vacuolar membrane transporter VIT1. Science 314, 1295–1298.

Krasileva, K. V. et al. (2017) Uncovering hidden variation in polyploid wheat. Proc. Natl. Acad. Sci., E913–E921.

Kushnirov, V. V. (2000) Rapid and reliable protein extraction from yeast. Yeast 16, 857–860.

Lamacchia, C., Shewry, P.R., Di Fonzo, N., Forsyth, J.L., Harris, N., Lazzeri, P.A., Napier, J.A., Halford, N.G., and Barcelo, P. (2001) Endosperm-specific activity of a storage protein gene promoter in transgenic wheat seed. J. Exp. Bot. 52, 243–250.

Lapinskas, Cunningham, K. W., Liu, X. F., Fink, G. R. and Culotta, V. C. (1995) Mutations in *PMR1* suppress oxidative damage in yeast cells lacking superoxide dismutase. Mol. Cell Biol. 15, 1382–1388.

Lapinskas, P.J., Lin, S.J., and Culotta, V.C. (1996) The role of the *Saccharomyces cerevisiae CCC1* gene in the homeostasis of manganese ions. Mol. Microbiol. 21, 519–28.

Lee, S., Jeon, U.S., Lee, S.J., Kim, Y.-K., Persson, D.P., Husted, S., Schjørring, J.K., Kakei, Y., Masuda, H., Nishizawa, N.K., and An, G. (2009) Iron fortification of rice seeds through activation of the nicotianamine synthase gene. Proc. Natl. Acad. Sci. U. S. A. 106, 22014–22019.

Li, L., Chen, O.S., McVey Ward, D., and Kaplan, J. (2001) CCC1 is a transporter that mediates vacuolar iron storage in yeast. J. Biol. Chem. 276, 29515–9.

MacDiarmid, C.W., Milanick, M.A., and Eide, D.J. (2003) Induction of the *ZRC1* metal tolerance gene in zinc-limited yeast confers resistance to zinc shock. J. Biol. Chem. 278, 15065–15072.

Moore, K.L., Zhao, F.-J., Gritsch, C.S., Tosi, P., Hawkesford, M.J., McGrath, S.P., Shewry, P.R., and Grovenor, C.R.M. (2012) Localisation of iron in wheat grain using high resolution secondary ion mass spectrometry. J. Cereal Sci. 55, 183–187.

Mumberg, D., Müller, R., and Funk, M. (1995) Yeast vectors for the controlled expression of heterologous proteins in different genetic backgrounds. Gene 156, 119–122.

Nakanishi, Y., Saijo, T., Wada, Y., and Maeshima, M. (2001) Mutagenic Analysis of Functional Residues in Putative Substrate-binding Site and Acidic Domains of Vacuolar H+-Pyrophosphatase. J. Biol. Chem. 276, 7654–7660.

Narayanan, N., Beyene, G., Chauhan, R.D., Gaitán-Solis, E., Grusak, M.A., Taylor, N., and Anderson, P. (2015) Overexpression of Arabidopsis *VIT1* increases accumulation of iron in cassava roots and stems. Plant Sci. 240, 170–181.

Neal, A.L., Geraki, K., Borg, S., Quinn, P., Mosselmans, J.F., Brinch-Pedersen, H., and Shewry, P.R. (2013) Iron and zinc complexation in wild-type and ferritin-expressing wheat grain: implications for mineral transport into developing grain. J. Biol. Inorg. Chem. 18, 557–70.

O’Dell, B.L., De Boland, A.R., and Koirtyohann, S.R. (1972) Distribution of phytate and nutritionally important elements among the morphological components of cereal grains. J. Agric. Food Chem. 20, 718–723.

Ravet, K., Touraine, B., Kim, S.A., Cellier, F., Thomine, S., Guerinot, M. Lou, Briat, J.F., and Gaymard, F. (2009) Post-translational regulation of *AtFER2* ferritin in response to intracellular iron trafficking during fruit development in Arabidopsis. Mol. Plant 2, 1095–1106.

Schroeder, J.I., Delhaize, E., Frommer, W.B., Guerinot, M. Lou, Harrison, M.J., Herrera-Estrella, L., Horie, T., Kochian, L. V, Munns, R., Nishizawa, N.K., Tsay, Y.-F., and Sanders, D. (2013) Using membrane transporters to improve crops for sustainable food production. Nature 497, 60–6.

Singh, S., Keller, B., Gruissem, W., and Bhullar, N. (2016) Rice *NICOTIANAMINE SYNTHASE* 2 expression improves dietary iron and zinc levels in wheat. Theor. Appl. Genet.130, 283–292.

Smedley, M.A. and Harwood, W.A. (2015) Gateway(r)-compatible plant transformation vectors. In Agrobacterium Protocols: Volume 1, Methods in Molecular Biology, K. Wang, ed (Springer: New York), pp. 1–15.

Trijatmiko, K.R. et al. (2016) Biofortified indica rice attains iron and zinc nutrition dietary targets in the field. Nat. Sci. Reports 6, 19792.

Vasconcelos, M.W., Gruissem, W., and Bhullar, N.K. (2017) Iron biofortification in the 21st century: setting realistic targets, overcoming obstacles, and new strategies for healthy nutrition. Curr. Opin. Biotechnol. 44, 8–15.

Velu, G., Ortiz-Monasterio, I., Cakmak, I., Hao, Y., and Singh, R.P. (2014) Biofortification strategies to increase grain zinc and iron concentrations in wheat. J. Cereal Sci. 59, 365–372.

Wu, H., Sparks, C., Amoah, B., and Jones, H.D. (2003) Factors influencing successful Agrobacterium-mediated genetic transformation of wheat. Plant Cell Rep. 21, 659–68.

Zhang, Y., Xu, Y.-H., Yi, H.-Y., and Gong, J.-M. (2012) Vacuolar membrane transporters OsVIT1 and OsVIT2 modulate iron translocation between flag leaves and seeds in rice. Plant J. 72, 400–10.

Zhao, F.J., Su, Y.H., Dunham, S.J., Rakszegi, M., Bedo, Z., McGrath, S.P., and Shewry, P.R. (2009) Variation in mineral micronutrient concentrations in grain of wheat lines of diverse origin. J. Cereal Sci. 49, 290–295.

Zimmermann, M.B. and Hurrell, R.F. (2007) Nutritional iron deficiency. Lancet 370, 511–520.

